# SARS-CoV-2 neutralizing human antibodies protect against lower respiratory tract disease in a hamster model

**DOI:** 10.1101/2020.08.24.264630

**Authors:** Bart L. Haagmans, Danny Noack, Nisreen M.A. Okba, Wentao Li, Chunyan Wang, Theo Bestebroer, Rory de Vries, Sander Herfst, Dennis de Meulder, Peter van Run, Mart M. Lamers, Bart Rijnders, Casper Rokx, Frank van Kuppeveld, Frank Grosveld, Dubravka Drabek, Corine GeurtsvanKessel, Marion Koopmans, Berend Jan Bosch, Thijs Kuiken, Barry Rockx

## Abstract

Effective clinical intervention strategies for COVID-19 are urgently needed. Although several clinical trials have evaluated the use of convalescent plasma containing virus-neutralizing antibodies, the effectiveness has not been proven. We show that hamsters treated with a high dose of human convalescent plasma or a monoclonal antibody were protected against weight loss showing reduced pneumonia and pulmonary virus replication compared to control animals. However, a ten-fold lower dose of convalescent plasma showed no protective effect. Thus, variable and relatively low levels of virus neutralizing antibodies in convalescent plasma may limit their use for effective antiviral therapy, favouring concentrated, purified (monoclonal) antibodies.

## INTRODUCTION

On 31 December 2019, the World Health Organization (WHO) was informed of a cluster of cases of pneumonia of unknown cause in Wuhan City, Hubei Province of China ^1^. Subsequently a novel coronavirus (SARS-CoV-2), was identified and as of August 11^th^, WHO reported more than 20 million cases of SARS-CoV-2 infection worldwide, with over 700,000 deaths.

SARS-CoV-2 infection is characterized by a range of symptoms, including fever, cough, dyspnea and myalgia ^2^. In severe cases, SARS-CoV-2 infection can be complicated by acute respiratory distress syndrome leading to respiratory insufficiency and multi-organ failure ^3^.

An effective treatment is a high priority as SARS-CoV-2 continues to circulate in many regions, and there is a risk of additional future waves of infection. To date, WHO reported at least 166 vaccine candidates being in different stages of development while other efforts include the development of neutralizing antibodies for prevention and/or treatment of SARS-CoV-2 infection. Early during the outbreak, the usefulness of convalescent plasma transfusion was considered for treatment of severe cases ^4^. Several large clinical trials have now been initiated to evaluate the efficacy and safety of convalescent plasma treatment of SARS-CoV-2 patients ^5^. Data on the outcomes of these trials have been limited and to date, preliminary results from only a few small cohorts and one randomized clinical trial have been published. Results from a randomized clinical trial did not show a benefit ^6,7^, while results from the small cohorts suggested clinical benefit but lacked controls for proper interpretation^8-11^. Although preclinical research indicated a limited protective effect of hamster serum when given to hamsters infected with SARS-CoV-2 early in the disease course ^12,13^, effects of human plasma have not been analyzed in this animal model. Importantly, data on the level of neutralizing antibodies that are required to provide a clinically meaningful protective effect are not available.

Apart from convalescent plasma, different human monoclonal antibodies (MAb) against SARS-CoV-2 have been identified and characterized, for prophylactic and therapeutic use. Most of these studies have shown efficient neutralization of SARS-CoV-2 *in vitro*, but few antibodies have been evaluated for their efficacy *in vivo*. We previously determined that MAb 47D11 efficiently neutralizes both SARS-CoV and SARS-CoV-2 *in vitro* ^14^. In the present study, we used this MAb and two doses of human convalescent plasma, differing almost ten-fold in neutralizing antibody concentration, to evaluate the efficacy of prophylactic antibody treatment in a hamster model of moderate to severe SARS-CoV-2 pneumonia.

## RESULTS

### Characteristics of neutralizing antibodies

We pooled 6 convalescent plasma samples from PCR-confirmed COVID-19 patients. The samples were selected based on a minimum neutralizing antibody titer of 1:1280 (PRNT_50_; **Supplementary Table 1**). The neutralizing antibody titer of the pooled plasma as well as the diluted pooled plasma were determined to be 1:2560 and 1:320 respectively (**Supplementary Table 1**). Only ten of 115 convalescent plasma donors previously tested had a titer of 1:2560 or higher while the 1:320 titer of the diluted plasma was just above the median titer of 1:160 of all donors tested ^7^.

In addition, we used human MAb 47D11 directed against SARS-CoV, which cross-reacts with SARS-CoV-2 and targets a conserved epitope in the S1 domain, previously shown to neutralize SARS-CoV-2 with an IC_50_ of 0.57 μg/ml ^14^. At a concentration of 3mg/mL the human MAb 47D11 preparation had an equivalent neutralizing antibody titer of 1:5260.

### Neutralizing antibodies protect against body weight loss from SARS-CoV-2 infection

To date, the Syrian golden hamster is the only animal species in which experimental SARS-CoV-2 infection results in moderate to severe pneumonia, with clinical signs, as well as shedding of virus ^12,13,15^. Therefore, the prophylactic potential of the 47D11 MAb and convalescent human plasma was evaluated in this hamster model. Twenty-four hours prior to challenge with SARS-CoV-2, animals were treated with MAb 47D11 or human convalescent plasma from COVID-19 patients. Volumes of human plasma treatment were chosen to mimic the application in humans. Animals were treated via intraperitoneal administration with either 3 mg MAb in 1mL (equivalent of a PRNT_50_ of 1:5260) or 500 μl human convalescent plasma (comparable to 300mL of convalescent plasma treatment in an adult human) containing either high (PRNT_50_ 1:2560) or median (PRNT_50_ 1:320) levels of SARS-CoV-2 neutralizing antibodies. Therefore, animals treated with human convalescent plasma were given a 4x or 40x lower dose of neutralizing antibodies compared to MAb 47D11, respectively. Unfortunately, due to technical restrictions blood could not be obtained on day 0 to determine the circulating neutralizing antibody titer. However, with a mean total blood volume in hamsters of 7,8 mL, we assumed that administration of MAb, and human convalescent plasma with high or median levels of neutralizing antibodies, lead to circulating neutralizing antibody titers of approximately 1:67, 1:16 and 1:2 respectively. There were three control groups, consisting of hamsters that were not treated prior to SARS-CoV-2 inoculation, and hamsters that were treated either with an irrelevant isotype control MAb or with normal healthy human plasma (not containing neutralizing antibodies to SARS-CoV-2; **Supplementary Table 1**) 24 hr before SARS-CoV-2 inoculation.

In line with earlier studies ^12,13^, experimental SARS-CoV-2 infection via the intranasal route resulted in a transient but significant weight loss in untreated animals as early as 3 days post inoculation (p.i.), approaching 20% weight loss by day 5 p.i. and normalizing by day 10 p.i. (**Figure 1A**). No other clinical signs were observed. Treatment with MAb 47D11 or a high dose of convalescent plasma protected animals against significant weight loss (**Figure 1A**). In contrast, treatment with the diluted convalescent plasma, control plasma, or control MAb did not protect against significant weight loss, with animals approaching 20% weight loss by day 5 p.i..

**Fig.1.**
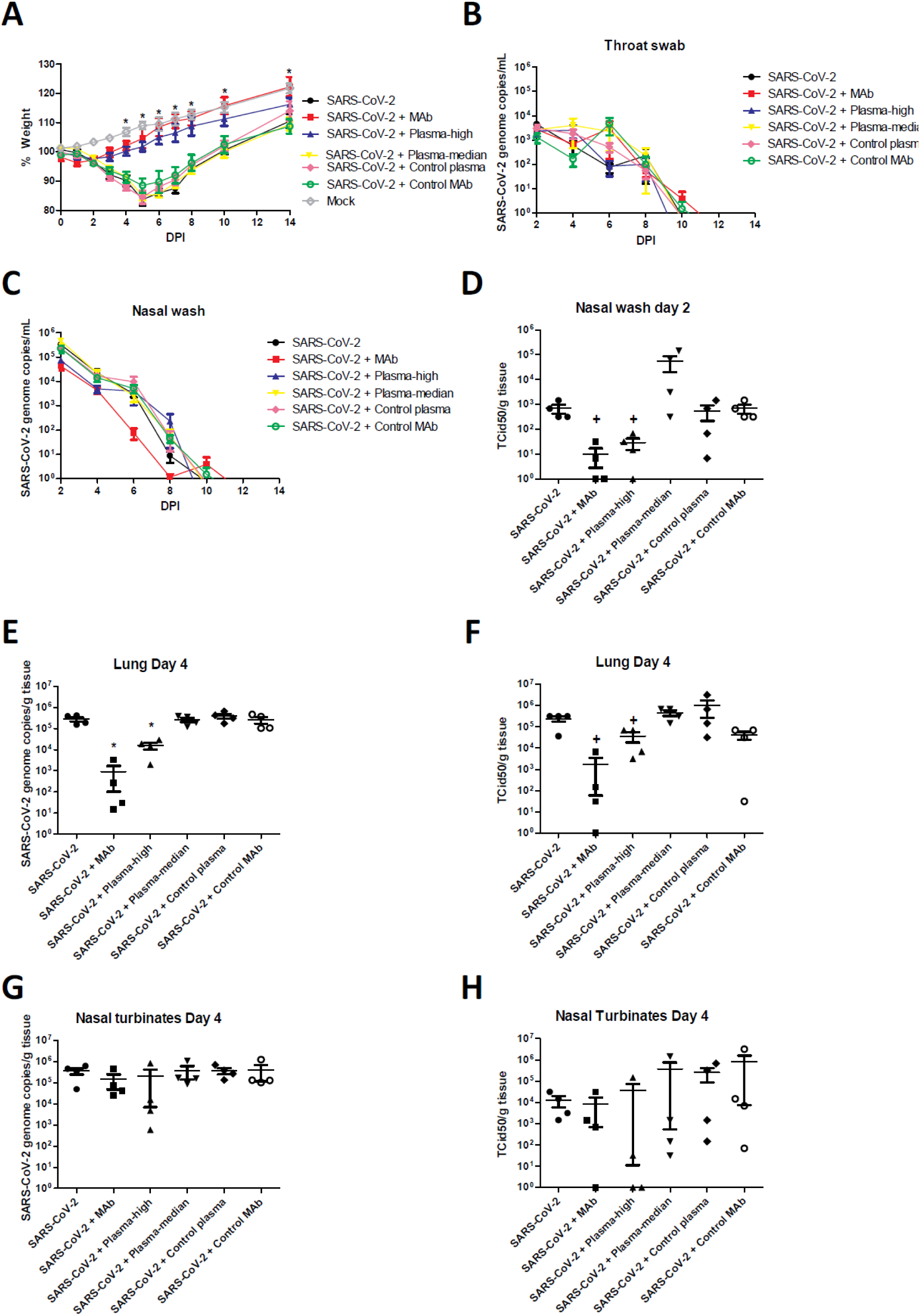
Effect of prophylactic neutralizing antibody treatment on weight loss and virus replication following SARS-CoV-2 infection in hamsters. A. Body weights of hamsters treated with antibodies were measured at indicated days after inoculation with SARS-CoV-2. SARS-CoV-2 viral RNA (B, C, E and G) or infectious virus (D, F and H) was detected in throat (B), nasal washes (C and D), lung (E and F) and nasal turbinates (G and H). The mean % of starting weight, the mean copy number or the mean infectious titer is shown, error bars represent the standard error of mean. n = 4. * = P<0.01 and + = P<0.05, ANOVA compared to SARS-CoV-2 inoculated, untreated animals.

### Minimal effect of antibody treatment on SARS-CoV-2 shedding

SARS-CoV-2 inoculation of hamsters resulted in detection of viral RNA in throat swabs from all groups for up to 10 days p.i., with peak shedding on day 2 p.i. (**Figure 1B)**. In addition, viral RNA was detected in nasal washes for up to 10 days p.i. (**Figure 1C**). While animals were protected against weight loss following treatment with either MAb 47D11 or high dose convalescent plasma, no significant reduction of viral RNA in throat swabs or nasal washes was observed. Despite high levels of viral RNA in nasal washes for several days p.i., infectious virus could only be isolated on day 2 p.i. (**Figure 1D**). Interestingly, while no significant effect of treatment was found on viral RNA detection, both treatment with MAb 47D11 and high dose convalescent plasma resulted in significant reduction of 1-2 logs in infectious virus on day 2 p.i. (p<0.05, ANOVA; **Figure 1D**).

Low levels of viral RNA were detected in rectal swabs on day 2 p.i. and occasionally at very low levels on other days in individual animals. There was no significant difference in virus detection in rectal swabs between treated and control groups and no infectious virus was detected.

### Antibody treatment reduced SARS-CoV-2 replication in the lower respiratory tract

Virus replication in the lungs and nasal turbinates was examined on day 4 p.i. (**Figure 1E-H**). In the lungs, treatment with MAb 47D11 or plasma with high neutralizing antibodies resulted in significant reduction of viral loads (both viral RNA, p<0.01 and infectious virus, p<0.05, ANOVA) (**Figure 1E and F**). In contrast, these treatments did not result in a significant reduction of viral load in the nasal turbinates (**Figure 1G and H**).

### Antibody treatment reduces histopathological changes in the respiratory tract following SARS-CoV-2 infection

At autopsy on day 4 p.i., control treated hamsters had single or multiple foci of pulmonary consolidation, visible as well-delimited, dark red areas, and covering 50-90% of the lung surface (**Figure 2**). No gross lesions were observed in any of the animals treated with either MAb 47D11 or high dose of convalescent plasma. Lungs from animals treated with the diluted plasma, or control plasma/ control MAb showed similar lesions to untreated animals.

**Fig. 2.**
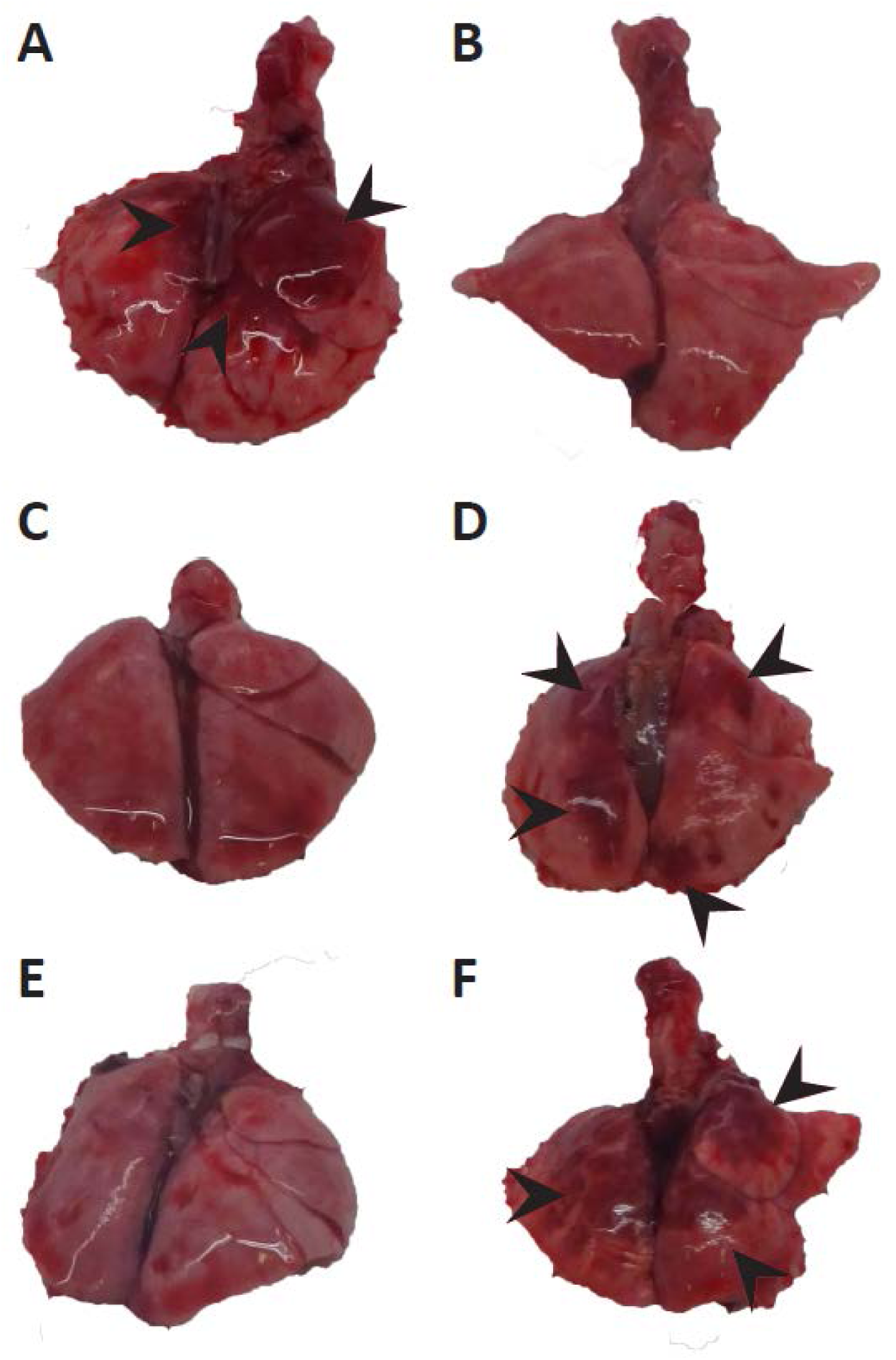
Gross pathological examination of the lungs of SARS-CoV-2 infected hamsters. Foci (arrowheads) of pulmonary consolidation in untreated SARS-CoV-2 infected animals (A) and animals treated with control MAb (D) or low dose plasma (F). Protection against pulmonary lesions in hamsters treated with MAb 47D11 (C) and high dose plasma (E), similar to mock infected animals (B). Images are from representative animals of each treatment group.

All animals, including the MAb 47D11 and high dose convalescent plasma groups, showed acute necrotizing and seropurulent rhinitis in the nasal cavity (**Figure 3**). It was centered on the olfactory mucosa, where it was marked and locally extensive. There, it was characterized by edema in the lumen mixed with sloughed epithelial cells, neutrophils, and cell debris, and by the presence of a moderate number of neutrophils in the epithelium and underlying lamina propria. Many cells in the olfactory epithelium in all animals expressed SARS-CoV-2 antigen, as demonstrated by immunohistochemistry. The inflammation in the mucosa and respiratory mucosa of the nasal cavity was mild and multifocal, and a few undifferentiated epithelial cells and ciliated columnar respiratory epithelial cells expressed virus antigen but did not differ between treated and untreated animals inoculated with SARS-CoV-2.

**Fig. 3.**
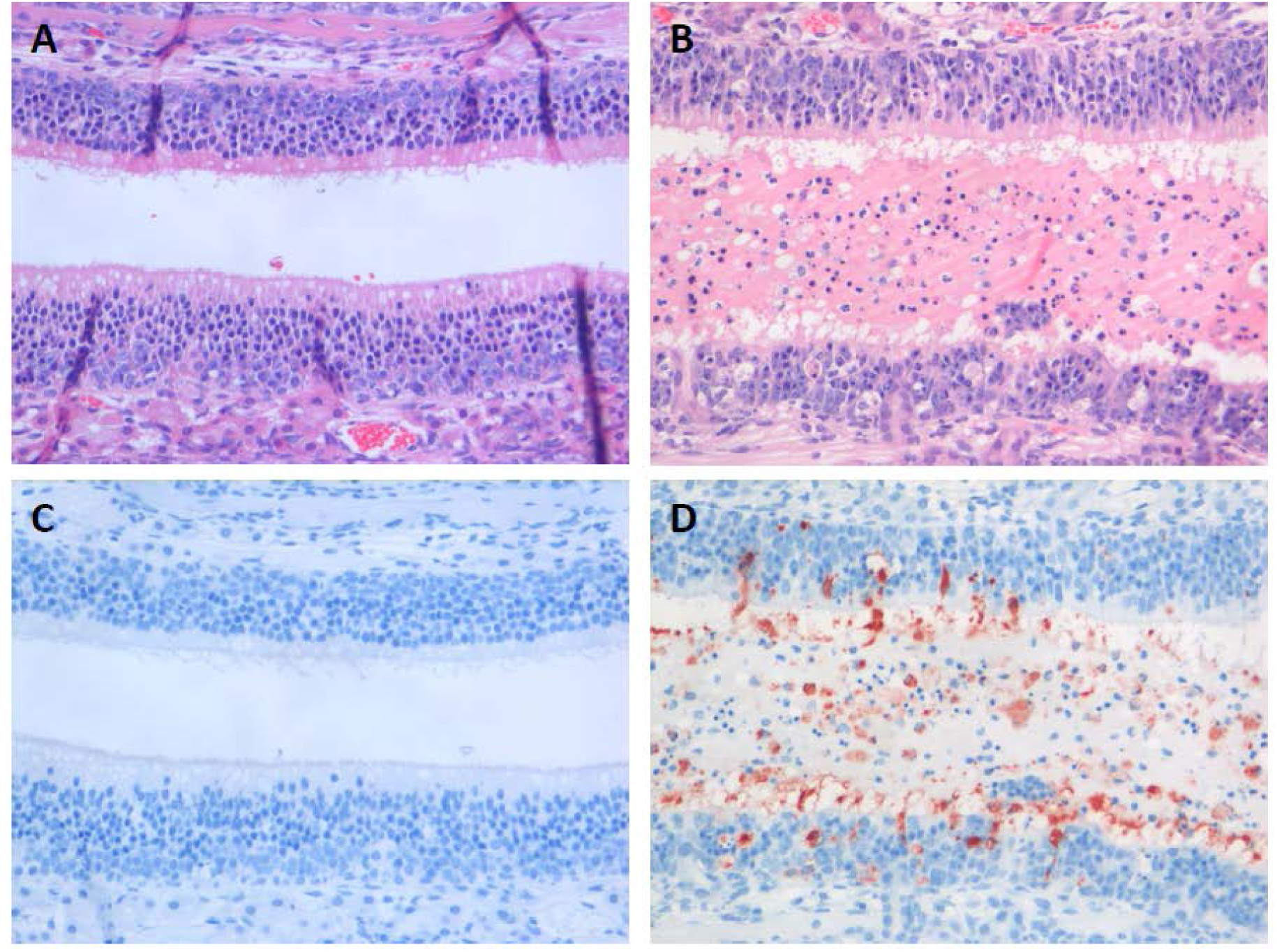
Histopathological changes and virus antigen expression in nasal turbinates of hamsters after challenge with SARS-CoV-2. In the nasal turbinate of a sham-inoculated hamster (left column), the nasal cavity is empty and the histology of the olfactory mucosa is normal (A). In a serial section, there is no SARS-CoV-2 antigen expression (C). In the nasal turbinate of a non-treated SARS-CoV-2-inoculated hamster (B and D), the nasal cavity is filled with edema fluid mixed with inflammatory cells and debris and the olfactory mucosa is infiltrated by neutrophils (B). A serial section of this tissue shows SARS-CoV-2 antigen expression in many olfactory mucosal cells, as well as in cells in the lumen (C).

The main observation in the lungs of the non-treated animals and the animals treated with diluted convalescent plasma, control plasma, or control MAb was multifocal or coalescing diffuse alveolar damage, which was characterized by loss of histological architecture of the lung parenchyma, edema, fibrin, sloughed epithelial cells, cell debris, neutrophils, mononuclear cells, and erythrocytes (**Extended Data Figure 1**). By immunohistochemistry, many type I pneumocytes and fewer type II pneumocytes at the edges of the lesions expressed virus antigen. Besides diffuse alveolar damage, there also was mild multifocal necrotizing and purulent bronchiolitis, characterized by loss of bronchiolar epithelium and the presence of a few neutrophils in the bronchiolar walls and lumina. By immunohistochemistry, a few bronchiolar epithelial cells expressed virus antigen.

Treatment with the 47D11 MAb resulted in a significant reduction of inflammation in the lungs (p<0.01, ANOVA; **Figure 4 and 5A**) and viral antigen expression in the lungs (p<0.05, ANOVA; **Figure 4 and 5B**). Although a reduction in inflammation and viral antigen was observed in the lungs of animals treated with high dose convalescent plasma, this was not statistically significant.

**Fig. 4.**
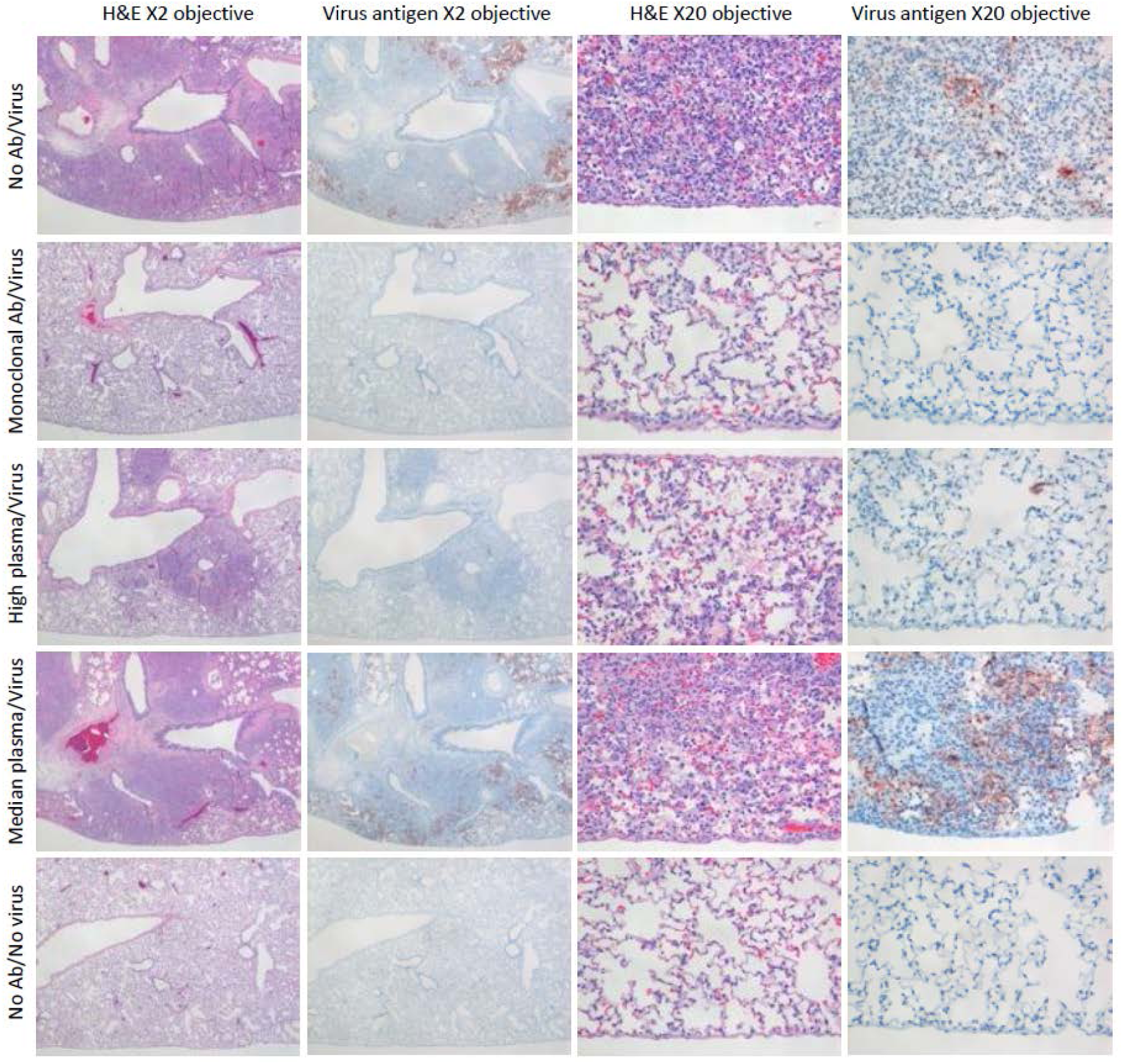
Effect of preventive treatment with MAb or high dose convalescent plasma on severity of pneumonia and level of virus antigen expression in lung parenchyma of hamsters after challenge with SARS-CoV-2. Comparison of extent of histopathological changes (HE) and virus antigen expression (IHC) at four days after SARS-CoV-2 inoculation at low magnification (two left columns) and high magnification (two right columns) in hamsters treated 24 hours before virus inoculation with neutralizing antibodies (second, third and fourth rows) compared to no treatment before SARS-CoV-2 inoculation (first row) and sham inoculation (fifth row).

**Fig. 5.**
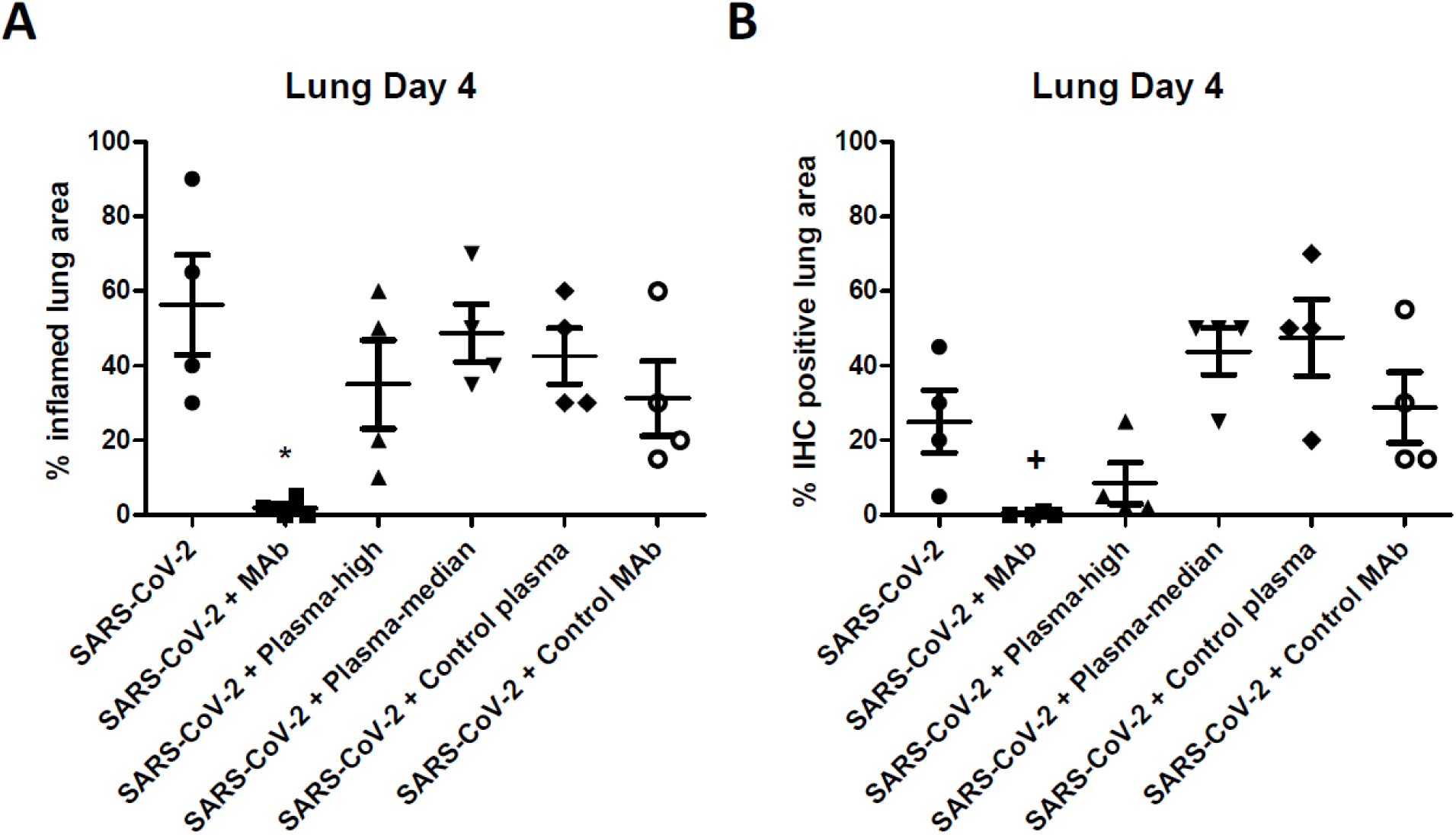
Quantitative assessment of histopathological changes and virus antigen expression. Percentage of inflamed lung tissue (A) and percentage of lung tissue expressing SARS-CoV-2 antigen (B) estimated by microscopic examination in different groups of hamsters at four days after SARS-CoV-2 inoculation. Individual (symbols) and mean (horizontal lines) percentages are shown. Error bars represent the standard error of mean. n = 4. * = P<0.01 and + = P<0.05, ANOVA compared to SARS-CoV-2 inoculated, untreated animals.

Following SARS-CoV-2 inoculation, all animals seroconverted by day 22 regardless of the treatment regimen (**Supplementary Table 2**). There was no significant difference in SARS-CoV-2 specific IgG titers among treatment groups with IgG titers of 1:12.800.

## DISCUSSION

Several studies have identified and characterized neutralizing antibodies against SARS-CoV-2 as a potential component of protective immunity ^14,16-22^. However, to date, only very few studies have focused on evaluating the efficacy of antibodies to protect or prevent against SARS-CoV-2 infection or disease *in vivo*. Those studies focused mainly on clinical signs and infection in the lungs and demonstrated mixed results with reduction in virus replication but no protection against pulmonary lesions ^13,16^, complicating the interpretation of data.

This study shows that prophylactic treatment with neutralizing antibodies prevents SARS-CoV-2 induced pneumonia in a hamster model. Animals treated with a high dose of neutralizing antibodies were protected against significant weight loss, did not show any gross lesions in their lungs and treatment resulted in a very substantial reduction in lung inflammation and virus replication in the lungs.

In agreement with recent studies, we show that prophylactic treatment with neutralizing antibodies can protect against disease following SARS-CoV-2 infection ^12,13^. While hamsters infected with SARS-CoV-2 showed no overt respiratory signs, they lose significant weight similar to what has been reported previously ^13,15^. Animals treated with high titers of neutralizing antibodies were protected against significant weight loss, did not show any gross lung lesions and had significantly less histological lesions and associated virus antigen expression in the lungs. Previous studies using convalescent hamster serum and a MAb showed that prophylactic treatment decreased virus replication in lungs similar to our findings, however the hamster serum did not protect against lung pathology ^12,13,15^. This is likely due to the fact that a lower dose of neutralizing antibodies was used (1:427) than was efficacious in this study (1 ml of MAb with titer 1:5260, or 0.5 ml of convalescent plasma with titer 1:2560).

Using convalescent plasma with lower neutralizing antibody titers (0.5 ml of convalescent plasma with titer 1:320), but still comparable to the median neutralizing titer found in patients recovered from COVID-19 ^7^, the protective efficacy was completely annulled. From our study the minimal protective neutralizing antibody titer in 0,5mL human plasma is between 1:320 and 1:2560. However, extrapolation to the human setting should be done with caution and studies on the levels and kinetics of neutralizing antibodies observed in humans after treatment with convalescent plasma are needed. In the current study we inoculated animals with a high dose of virus and by a method that, despite intranasal inoculation, ensures delivery of virus in the lower respiratory tract. While this results in a robust model of SARS-CoV-2 pneumonia, humans will most likely be exposed to a much lower level of virus. Nevertheless, these data highlight the importance of pre-screening convalescent plasma from donors prior to use for convalescent plasma treatment. Indeed, levels of neutralizing antibodies vary substantially between individuals with a recent study showing a median titer of 1:160 in convalescent plasma in 115 donors and 22% had a titer of 1:40 or lower ^7^. The lower titers are more typically observed after mild or asymptomatic COVID-19 cases ^23^; those that actually act as plasma donor.

While prophylactic treatment resulted in protection against disease and reduced SARS-CoV-2 replication in the lungs, only a limited effect was found in the upper respiratory tract. Previous studies with influenza virus have shown that serum IgG can diffuse into alveolar lining fluid, thus protecting the lung parenchyma against virus infection ^24^. In contrast, the concentration of IgG on the surface of nasal mucosa is much lower. This suggests that treatment may protect against disease in the lungs but not virus transmission from the nose. Recent studies have shown that SARS-CoV-2 can transmit between animals via both direct contact and air ^13,15,25^. Similar to our study, infectious virus was only detected in nasal washes early during infection and the period in which virus could be transmitted to naïve animals correlated with the presence of infectious virus ^15^. All animals treated with the MAb and convalescent plasma seroconverted, therefore, antibody based prevention of COVID-19 did not seem to prevent the development of humoral immunity after SARS-COV-2 exposure.

To date, the efficacy of prophylactic antibody treatment has not been evaluated in humans. Several studies have reported on the possible efficacy and safety of therapeutic treatment with convalescent plasma in both small cohorts as well as a clinical trial, with variable and inconclusive results ^6,9-11^. The main results from the small cohorts suggest a clinical benefit and reduced virus loads; however, given the limited information and lack of controls, interpretation of the data is inconclusive. Recently, two randomized clinical trials were prematurely terminated and did not result in a shorter time to clinical improvement ^6,7^. The effect of treatment may be limited due to use of convalescent plasma with low levels of neutralizing antibodies of at least 1:40 to 1:80. Given the dilution factor upon intravenous administration, this would effectively result in neutralizing antibody titers of ∼1:2 or 1:4, respectively.

Unfortunately, in the current study, we were not able collect blood at the time of virus challenge. Furthermore, a recent study showed that most COVID-19 patients already have neutralizing antibody titers of 1:60 or higher at hospital admission ^7^, supporting our findings that only treatment with high levels of neutralizing antibodies may have a protective effect. In addition, in most studies, only severe cases of SARS-CoV-2 infection were included at the time when patients were admitted to a hospital for severe disease. At that time, the therapeutic window for antibody treatment may have passed since many patients with severe disease are already resolving the virus infection in the lung while the observed severe disease is primarily due to an aberrant host response rather than virus infection. Hence for effective treatment, the timing and dosing of administered neutralizing antibodies is likely critical.

Despite being promising for prevention and treatment of COVID-19 infection, the use of hyperimmune globulin preparations from recovered patients has its inherent challenges, including safety, batch-to-batch variation, scalability, standardized dosing and presence of non-neutralizing antibodies. In addition, the potential for antibody dependent enhancement (ADE) of disease remains a concern in the development of vaccines and antibody treatments ^26^. Experimental studies have previously shown that antibodies against SARS-CoV can induce severe lung injury ^27^. In the current study we did observe increased levels of infectious virus in the nasal washes of some individual animals treated with lower levels of neutralizing antibodies, but not significant due to the variation in infectious titers between animals, and was also not supported by significant difference in disease or histopathology. Given that we cannot reliably predict ADE of disease after antibody treatment or vaccination, it will be crucial to evaluate safety in humans as convalescent plasma treatments continue. These challenges are more easily addressed by using purified and concentrated plasma derived antibodies or (combinations) of recombinantly produced MAbs. MAbs with desired properties can be selected from the immune repertoire of e.g. infected or immunized individuals with respect to binding affinity, potency and breath of neutralization. Moreover, antibody engineering allows to tweak the Fc-mediated immune effector functions and to improve MAb pharmacokinetics and reduce potential disease enhancing effects. In addition, established manufacturing pipelines allow efficient, highly controlled and scalable production. Success with single or combinations of monoclonal antibodies has recently been achieved for treatment of Ebolavirus ^28^, whereas other treatments, including convalescent plasma, did not show a benefit ^29^.

In conclusion, our data show that prophylactic treatment with a highly neutralizing MAb not only protects against weight loss and reduces virus replication in the lungs, it also limits histopathological changes in the lungs. In addition, we show that while prophylactic treatment may prevent disease, animals still become infected and shed virus, indicating that transmission will not be blocked. These data highlight the importance to include virus shedding, replication in lungs as well as clinical and pathological determinants of disease in evaluating the efficacy of antibody treatment. In contrast, treatment with convalescent plasma provides only partial protection, and only when plasma with high neutralizing titers was used. This protective effect is completely annulled when using the median neutralizing antibody dose found in recovered patients ^7^. It is therefore crucial to select convalescent plasma from donors with high levels of neutralizing antibody. Given the variation in antibody responses in patients, this limits the number of suitable donors for preparing immunoglobulin therapies considerably. No such limitation is present with *in vitro* produced MAbs and our results suggest this may be the more favorable route to develop an effective therapy.

## MATERIALS AND METHODS

### Viruses and cells

SARS-CoV-2 (isolate BetaCoV/Munich/BavPat1/2020) was obtained from a clinical case in Germany diagnosed after returning from China (European Virus Archive Global # 026V-03883). The virus was propagated to passage three on Vero E6 cells in Opti-MEM I (1X) + GlutaMAX (Gibco), supplemented with penicillin (10,000 IU/mL) and streptomycin (10,000 IU/mL) at 37°C in a humidified CO2 incubator. All work was performed in a Class II Biosafety Cabinet under BSL-3 conditions at the Erasmus Medical Center (MC).

### MAbs and convalescent plasma

We previously identified MAb 47D11 which efficiently neutralizes SARS-CoV-2 *in vitro* ^14^. The irrelevant isotype control antibody used in this study was characterized previously^32^.

Convalescent plasma was collected from donors who had a RT-PCR confirmed SARS-CoV-2 infection and were asymptomatic for at least 14 days ^7^. Of all donors tested, only plasma with neutralizing antibodies against SARS-CoV-2 confirmed by a SARS-CoV-2 plaque reduction neutralization test (PRNT) and a PRNT_50_ titer of at least 1:1280 was used. Equal volumes of plasma from 6 donors was pooled and used for prophylactic treatment in hamsters (High dose). In addition, the pooled plasma was diluted 10-fold in PBS (Median dose). Normal human plasma from a healthy donor was used as a control.

### Animals and Ethical Statement

Animals were handled in an ABSL3 biocontainment laboratory. Research was conducted in compliance with the Dutch legislation for the protection of animals used for scientific purposes (2014, implementing EU Directive 2010/63) and other relevant regulations. The licensed establishment where this research was conducted (Erasmus MC) has an approved OLAW Assurance # A5051-01. Research was conducted under a project license from the Dutch competent authority and the study protocol (#17-4312) was approved by the institutional Animal Welfare Body. Animals were housed in groups of 2 animals in filter top cages (T3, Techniplast), in Class III isolators allowing social interactions, under controlled conditions of humidity, temperature and light (12-hour light/12-hour dark cycles). Food and water were available ad libitum. Animals were cared for and monitored (pre- and post-infection) by qualified personnel. The animals were sedated/anesthetized for all invasive procedures.

### Animal procedures SARS-CoV-2

Female Syrian golden hamsters (*Mesocricetus auratus*; 6-week-old hamsters from Janvier, France) were anesthetized by chamber induction (5 liters 100% O2/min and 3 to 5% isoflurane). 24-hour prior to inoculation with virus, groups of 8 animals were treated with either 3mg of MAb in 1mL or 500 μl human convalescent plasma via the intraperitoneal route.

Animals were inoculated with 10^5^ TCID50 of SARS-CoV-2 or PBS (mock controls) in a 100 μl volume via the intranasal route. During the experiment the animals were monitored for general health status and behavior daily and were weighed regularly for the duration of the study (up to 22 days post inoculation; d.p.i.). Nasal washes, throat swabs and rectal swabs were collected under isoflurane anesthesia during the study. Groups of 4 animals were euthanized on day 4 or day 22 after inoculation, and serum samples, as well as lung, and nasal turbinates, were removed for virus detection and histopathology.

### Serological Analysis

To test for SARS-CoV-2 antibodies, hamster serum samples were collected at days 4 and 22. Serum samples were tested for SARS-CoV-2 antibodies using a spike S1 and nucleocapsid protein (N) ELISAs ^23^. Briefly, ELISA plates were coated overnight with SARS-CoV-2 S1. After blocking, serum samples were added and incubated for 1h at 37°C. Bound antibodies were detected using HRP-labelled rabbit anti–human IgG (Dako) or anti-hamster IgG and TMB (Life Technologies) as a substrate. The absorbance of each sample was measured at 450 nm.

A plaque reduction neutralization test (PRNT) was used as a reference for this study as previously described ^23^. The serum neutralization titer is the reciprocal of the highest dilution resulting in an infection reduction of >50% (PRNT50).

### Virus detection

Samples from nasal turbinates and lungs were collected post mortem for virus detection by RT-qPCR and virus isolation as previously described ^30^. Briefly, tissues were homogenized 10% w/v in viral transport medium using Polytron PT2100 tissue grinders (Kinematica). After low-speed centrifugation, the homogenates were frozen at −70°C until they were inoculated on Vero E6 cell cultures in 10-fold serial dilutions. The SARS-CoV-2 RT-qPCR was performed and quantified as copy numbers as previously published ^31^.

### Histopathology and immunohistochemistry

For histological examination lung and nasal turbinates were collected. Tissues for light-microscope examination were fixed in 10% neutral-buffered formalin, embedded in paraffin, and 3 μm sections were stained with haematoxylin and eosin.

Sections of all tissue samples were examined for SARS-CoV-2 antigen expression by immunohistochemistry as previously described ^30^. Briefly, paraffin was removed from sections, and viral antigen was detected using a rabbit polyclonal antibody against SARS-CoV-nucleoprotein (40143-T62, Sino Biological, Chesterbrook, PA, USA) and horseradish peroxidase labeled goat-anti-rabbit IgG (P0448, DAKO, Agilent Technologies Netherlands B.V. Amstelveen, The Netherlands). Horseradish peroxidase activity was revealed by incubating slides in 3-amino-9-ethylcarbazole (Sigma, St Louis, MO, USA) solution, resulting in a bright red precipitate. Sections were counterstained with haematoxylin.

For quantitative assessment of SARS-CoV-2 infection-associated inflammation in the lung, each H&E-stained section was examined for inflammation by light microscopy using a 2.5x objective, and the area of visibly inflamed tissue as a percentage of the total area of the lung section was estimated. Quantitative assessment of virus antigen expression in the lung was performed according to the same method, but using lung sections stained by immunohistochemistry for SARS-CoV-2 antigen. Sections were examined without knowledge of the identity of the hamsters.

### Statistical analysis

Statistical analyses were performed using GraphPad Prism 5 software (La Jolla, CA, USA). Each specific test is indicated in the figure legends. P values of ≤0.05 were considered significant. All data are presented as means ± standard error of the mean (SEM).

## ACKNOWLEDGEMENTS

We thank J.M. Fentener van Vlissingen, Y. Kap, D. Akkermans, V. Vaes, for assistance with the animal studies. This research is (partly) financed by the NWO Stevin Prize awarded to M.K. by the Netherlands Organisation for Scientific Research (NWO), H2020 grant agreement 874735-VEO to M.K., the Netherlands Organisation for Health Research and Development (ZONMW) grant agreement 10150062010008 to B.H. and H2020 grant agreement 101003651–MANCO to B.H. The research was co-funded by the PPP Allowance (grant agreement LSHM19136) made available by Health Holland, Top Sector Life Sciences & Health, to stimulate public-private partnerships.

## AUTHOR CONTRIBUTIONS

Conceptualization, B.R., M.K., B.H.; investigation, B.R., D.N., N.O., W.L., C.W., T.B., R.d.V., S.H., D.d.M., P.v.R., M.L.; resources, B.H., B.R., C.R., F.v.K., F.G., D.D., C.G.v.K., B.B.; supervision, B.R. and B.H.; writing, original draft, B.R., T.K., and B.H.; writing–review and editing, all authors; funding acquisition: B.H., F.G., B.B., and M.K.

