## Supplemental material for "SARS-CoV-2 neutralizing human antibodies protect against lower respiratory tract disease in a hamster model"

**Supplementary Table 1. SARS-CoV-2 neutralizing antibody titers in individual and pooled donor plasma.**

| | **$PRNT_{50}$ TITER** |
|---|---|
| **DONOR 1** | 1280 |
| **DONOR 2** | 1280 |
| **DONOR 3** | 1280 |
| **DONOR 4** | >2560 |
| **DONOR 5** | 1280 |
| **DONOR 5** | 2560 |
| **POOLED HIGH** | 2560 |
| **POOLED MEDIAN** | 320 |
| **POOLED NEGATIVE** | <20 |

**Supplementary Table 2. Neutralizing antibody response ($PRNT_{50}$) in hamsters following prophylactic treatment.**

| ANIMAL | SARS-COV-2 ONLY | MAB | PLASMA-HIGH | PLASMA-LOW | CONTROL PLASMA | CONTROL MAB | MOCK |
|---|---|---|---|---|---|---|---|
| **1** | 2560 | 2560 | 2560 | 2560 | 2560 | 2560 | <20 |
| **2** | 2560 | 2560 | 1280 | 2560 | 2560 | 640 | <20 |
| **3** | 2560 | 1280 | 2560 | 1280 | 2560 | 1280 | <20 |
| **4** | 2560 | 2560 | 640 | 2560 | 2560 | 1280 | <20 |

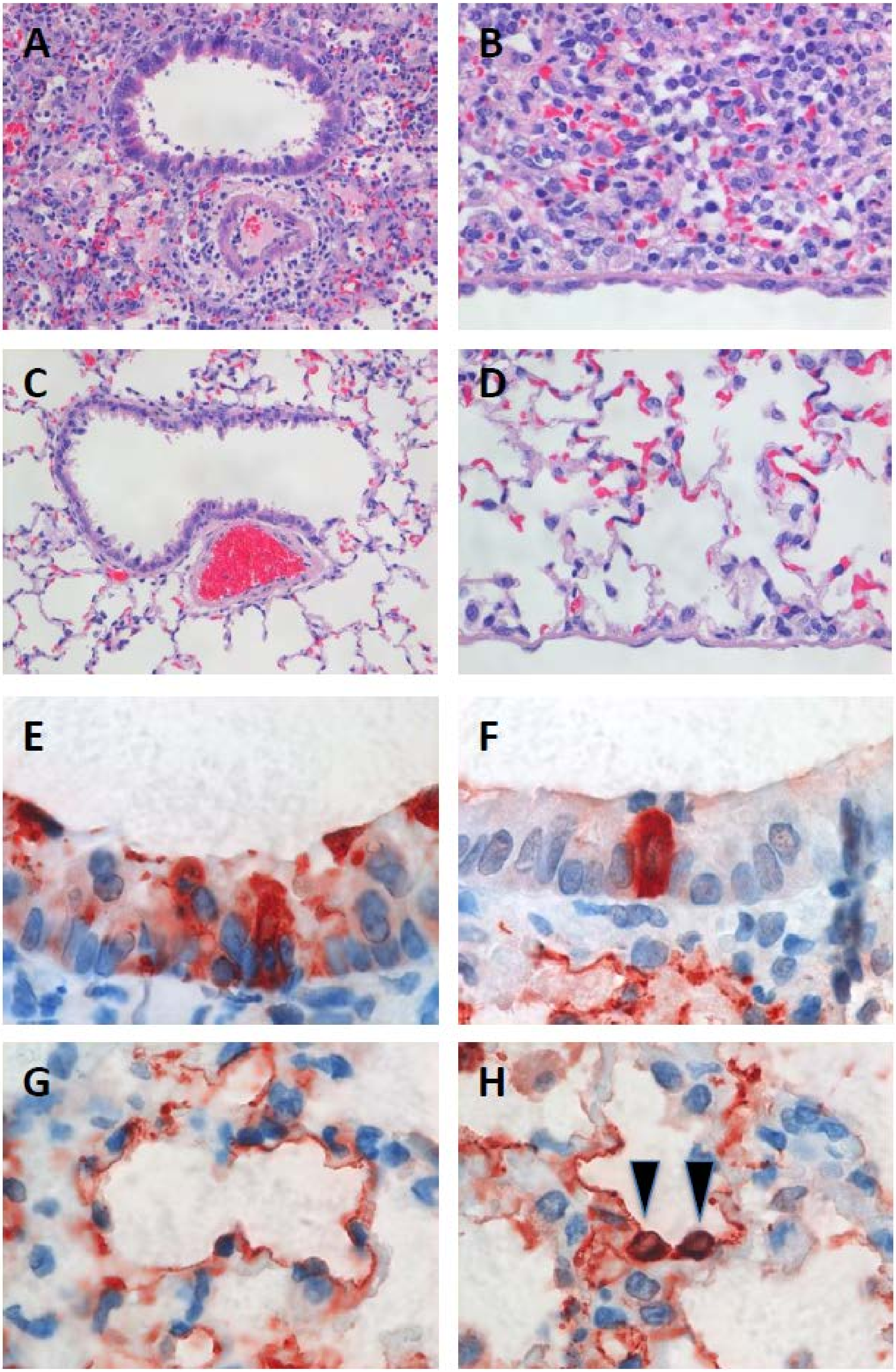
A
B
C
D
E
F
G
H

**Supplemental Fig. 1** Histopathological changes and virus antigen expression in bronchioles and alveoli of hamsters after challenge with SARS-CoV-2. In the lung of a non-treated SARS-CoV-2-inoculated hamster (A and B), the alveoli are flooded by edema fluid (B), fibrin, sloughed epithelial cells, cell debris, neutrophils, mononuclear cells, and erythrocytes, obfuscating the histological architecture, and there are a few inflammatory cells in the lumen and epithelium of a bronchiole (A), and in the perivascular space around an adjacent blood vessel. In the lung of a sham-inoculated hamster (C and D), the lumina of the alveoli are empty (D), the alveolar walls are thin, and there are no inflammatory cells in or around the walls of the bronchiole and adjacent blood vessel (C). The lung of a non-treated SARS-CoV-2-inoculated hamster shows SARS-CoV-2 antigen expression in multiple (E) or solitary (F) bronchiolar epithelial cells, type I pneumocytes (G), and type II pneumocytes (H, arrowheads).
